# Mosquito vector ecologies are destabilizing as a result of climate change

**DOI:** 10.1101/2025.09.15.676177

**Authors:** Evan J. Curcio, Kai Xu, Harutyun Sahakyan, Yuri I. Wolf, Elizabeth A. Kelvin, Nash D. Rochman

## Abstract

Mosquito-borne infectious disease is a major cause of mortality and a significant economic burden worldwide. Shifting regional and seasonal patterns make proactive intervention challenging. Here we introduce a machine learning approach trained on satellite and mosquito observational data, improving generalizability to sparse observations while retaining similar positive performance characteristics of models used in standard practice. We provide global estimates for *Aedes* habitability at high spatial resolution in five-year increments from 1970-2024. The vast majority of ecologies appear to be destabilizing. Overall, we demonstrate a decrease in risk near the equator and an increase in risk in historically temperate climates including the United States, the European Union, and China. Despite this decrease in risk relative to historical averages, regions near the equator remain among the highest risk worldwide. Together, these results reflect an overall, marked expanse of the regions of the globe that support *Aedes* mosquitos and we observe an ongoing, linear increase in the global population at risk of contracting mosquito-borne disease.

## 1. Introduction

Vector-borne disease accounts for about 17% of all infectious diseases, causing over 700,000 deaths annually. Mosquito-borne pathogens are responsible for hundreds of millions of human infections per year including malaria, dengue, lymphatic filariasis, chikungunya, West Nile fever, and Zika virus disease, among others (1). Nearly half of the global population is at risk of vector-borne disease (2–5), imposing significant economic costs stemming from government-sponsored pharmaceutical treatment and public health intervention; decreased economic output due to illness and private payments made by patients; and decreased revenues from tourism (6). Mosquito breeding cycles, biting frequency, lifespans, and habitat - as well as transmission characteristics for the viral and bacterial pathogens they carry - are affected by climatic variables including temperature, humidity, and precipitation (7–11). Consequently, climate change is altering the geographic range over which mosquito vectors are commonly found (12–15). Incorporating climate data into predictions for mosquito habitability may lead to more efficient and effective mitigation efforts.

A variety of mathematical modeling techniques have been explored, largely falling into two categories: mechanistic modeling, primarily supported by dynamical systems representing expert knowledge about vector biology, and statistical physics based approaches, primarily supported by detailed environmental and observational data (16–20). More recently, machine-learning approaches have shown promise towards making more generalizable predictions with reduced computational requirements, supporting improved speed, customizability, and scalability (21–24). Here, we use a simple feedforward neural network (FFNN) architecture trained on remote sensing data from the ERA5 project (25,26) and mosquito observations from iNaturalist (27) via a presence/pseudo-absence approach. We first characterize our approach, motivating that it provides improved generalizability to sparse observations while retaining similar positive performance characteristics of models used in standard practice, and proceed to quantify the impact of climate change on the ecology of three mosquito species: *Aedes aegypti, albopictus*, and *vexans*.

## 2. Results

### 2.1 Model performance

The data pipeline begins with the ERA5 climate data, downloaded in bulk (or optionally, piecewise through Earth Engine API) (28). Terrestrial air temperature range, median air temperature, relative humidity, total precipitation, and wind speed were computed daily at a resolution of 0.25 by 0.25 degrees. True positive locations where mosquitos have been observed were obtained from iNaturalist and pseudo-absences were drawn roughly uniformly from remaining terrestrial locations over the time window of iNaturalist observations, *A. aegypti*: Mar 2008 - Aug 2024; *albopictus*: Apr 2002 - Dec 2024; *vexans*: Sep 2003 - Jan 2025 (Figure 1A). Climate data and true positive / pseudo absence labels were then used to train a standard FFNN architecture ending in a binary cross entropy with logits loss layer yielding a value between 0 and 1 (Figure 1B). The distribution of losses is highly dependent on model hyperparameters including the true positive to pseudo absence ratio. To enable direct comparison among trained models, losses were converted to the corresponding false negative rates (FNR) for true positive locations with the same loss. The FNR may be interpreted as “risk percentiles”. For example, observers in date-locations with an FNR of 0.5 (50%) have a greater likelihood of observing a mosquito than they would if they were present at 50% of date-locations where observations were recorded.

**Figure 1:**
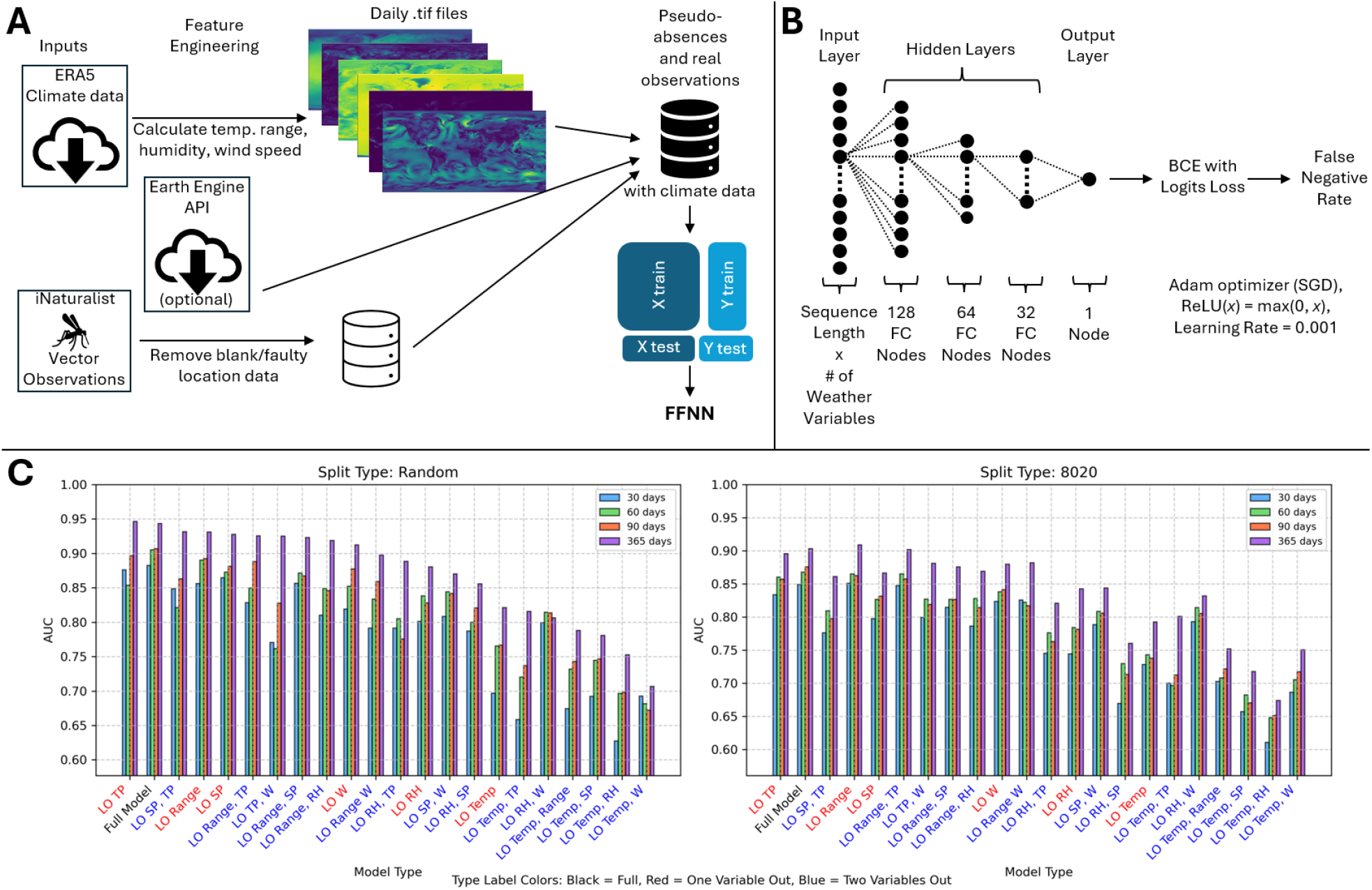
Model characterization. **A**. Data pipeline schematic. **B**. Feedforward neural network architecture. **C**. Leave-one-out and Leave-two-out analysis for *A. aegypti* in 2024. Sequence lengths (SL) tested included 30 days (blue bars), 60 days (green bars), 90 days (red bars), and 365 days (purple bars). Left: 80% train/20% test groups chosen randomly. Right: 80% train/20% test groups chosen chronologically (training using the oldest 80% of observations). Models (x-axis) ordered by highest AUC of all SLs. Model label color signifies full model (black), leave-one-out (red), or leave-two-out (blue). AUCs are computed according to the sensitivity/specificity curve varying the FNR threshold for positive labelling.

We evaluated model generalizability by chronologically separating training/test splits, reducing the duration of training data to as little as 30 days prior to the prediction day, and removing up to two of the four computed climate variables. We found model performance was insensitive to small perturbations. The relative performance of models trained using shorter durations was more variable than models trained using longer durations under variable reduction, as would be expected from overfitting or sensitivity to short-term weather pattern variation. The full model, including only climate variables widely understood to impact vector habitability (29–31), ranked second in both training/test splits (the models ranked first differed between training/test splits) and was selected for implementation (Figure 1C). See Methods for details.

### 2.2 Comparison with MaxEnt

We compared predictions from our FFNN approach to those obtained using MaxEnt, a maximum entropy modeling software (17). Per our review, MaxEnt remains the most commonly used program for species geospatial range prediction (19). MaxEnt is memory-intensive and requires specific input data formatting. Here we demonstrate that our approach is able to achieve qualitatively similar results to MaxEnt with a more flexible, lightweight pipeline with compute times on the order of a few minutes per daily prediction using 90 day sequence length with 16GB RAM. We also motivate that our approach provides improved generalization when labelled data are sparse. The results shown in Figure 2 utilize monthly WorldClim climate data (minimum temperature, maximum temperature, and precipitation at each grid point) from 2020 to predict January 2021 *A. aegypti* habitability, adopting the most commonly used input source for MaxEnt, for both models as well as ERA5 for our approach. Figure 2A identifies regions representing areas where vector-borne infections are of significant historical or increasing concern, due to changing climate, high human population, or increased human migration (32–41).

**Figure 2:**
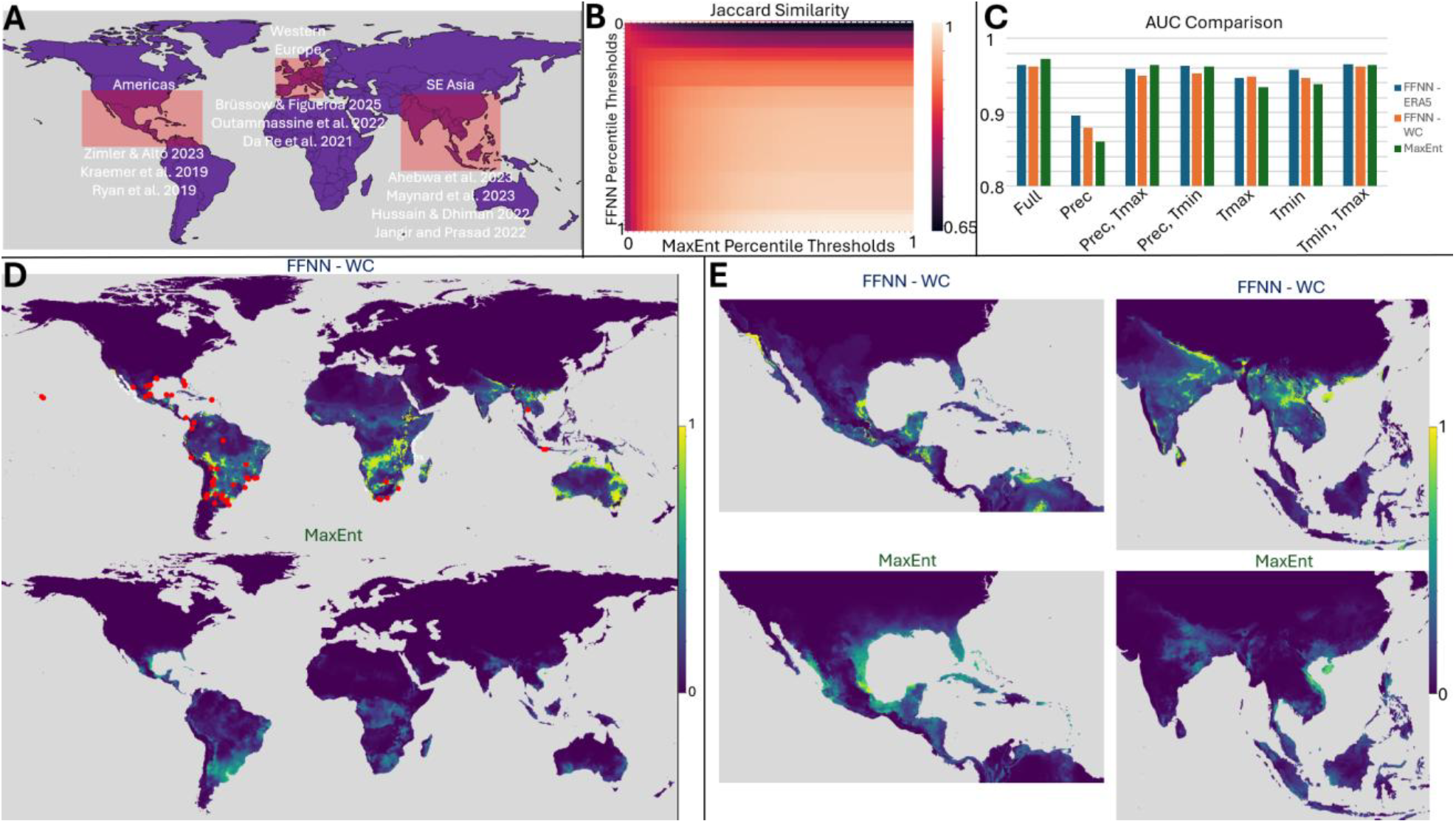
Comparison of the FFNN to MaxEnt using 2020 WorldClim data to predict Jan 2021 *A. aegypti* presence. **A**. Regions of study corresponding to areas of concern identified in literature (red rectangles, top left). **B**. Binary Comparison matrix (top middle), demonstrating Jaccard similarity between FFNN and MaxEnt. Difference in assignment of 0 probability is demonstrated by the top row (dashed white line). **C**. Histogram comparing AUC values for January 2021 predictions using monthly FFNN model trained on ERA5 data (blue), monthly FFNN model trained on WorldClim data (orange), and MaxEnt model trained on WorldClim data (green). **D**: Jan 2021 global *A. aegypti* false negative rates using WorldClim data with FFNN (top), shown with observation data (red markers) and MaxEnt (bottom). **E**: Comparison of *A. aegypti* false negative rates in the Americas (left) and SE Asia (right) for Jan 2021 using FFNN (top) and MaxEnt (bottom) trained on WorldClim data, demonstrating marked difference in predictions for Jan 2021.

Globally, our approach yields qualitatively similar predictions to MaxEnt across a wide range of sensitivities. Figure 2B displays the Jaccard similarity for binary predictions assigned on the basis of an FNR exceeding the specified threshold (note MaxEnt natively reports FNR as “cumulative” output). Leave-one-out analysis demonstrates overall superior performance within our model trained on ERA5 data over WorldClim data. Notably, leaving out precipitation had the largest negative impact on AUC values despite this model yielding the globally optimal score for one of the train/test splits presented in Figure 1. Regardless of the input data, however, our model yields slightly reduced AUCs in comparison to MaxEnt (Figure 2C).

We motivate that this is a modest cost for the improved generalizability to sparse label data the FFNN affords. Figure 2D contrasts global FFNN (top) and MaxEnt (bottom) predictions. Our approach assigns risk scores near 1 for a substantial portion of the globe in comparison to MaxEnt for which areas of high risk are tightly localized, primarily around true positive labels. Figure 2E focusing on regions of concern further highlights this contrast. Near the equator within the Western hemisphere, MaxEnt assigns a significant region high risk scores, qualitatively corresponding to our own, although our model assigns substantially higher risk to Baja California, the southwestern United States, the Yucatan peninsula, and much of Colombia and Venezuela. In fact, our model corroborates well-documented invasions of *A. aegypti* and *A. albopictus* in southern California (42). In the Eastern hemisphere, however, for which there are relatively fewer true positive observations (Figure 2D, red markers), MaxEnt predictions are significantly more conservative. This likely reflects underprediction given that the landmass suitable for mosquito habitability in the Eastern hemisphere is disproportionately undersampled relative to the Western hemisphere.

To summarize, our approach 1) produces qualitatively similar results to the widely used tool, MaxEnt, replicating regional relative risks where observational data is robust 2) shows the capacity to generalize well under conditions where true positive labels are sparse, predicting significantly higher risk across much of the globe and 3) supports the use of a greater diversity of input training data types with reduced memory requirements. For example, users may work with ERA5 as we do in this study or ERA5-Land (25,26). Both projects offer climatic variables at higher resolution than other publicly available datasets and prior work has shown that including ERA5 data in reanalysis can improve accuracy across regions of importance (43–49). We proceed to establish seasonal trends and historical changes in *Aedes* habitability in the following sections using our approach.

### 2.3 Seasonal trends

To account for yearly variation in short term weather patterns, daily FNR values (see Movies in Supplementary Data) at every location were averaged across January 1, 2020 to December 31, 2024 for each of the three species studied (Figure 3A). Averages were also taken of each season, which we identify as March through May (Spring), June through August (Summer), September through November (Fall/Autumn), and December through February (Winter). We then calculated differences between successive seasons, reflecting the intensity of seasonal transitions (Figure 3B). Spring to Summer represents the largest increase in the Northern hemisphere and largest decrease in the Southern hemisphere, while Fall to Winter represents the inverse (as expected). Overall, seasonal transitions are modestly more well defined in the Northern hemisphere. There are also significant species- and region-specific seasonal patterns. For example, the Indian subcontinent experiences seasonality characteristic of the Southern hemisphere for *A. vexans* but the Northern hemisphere for *A. aegypti* and *A. albopictus*, with implications for the optimal timing of mitigation strategies.

**Figure 3:**
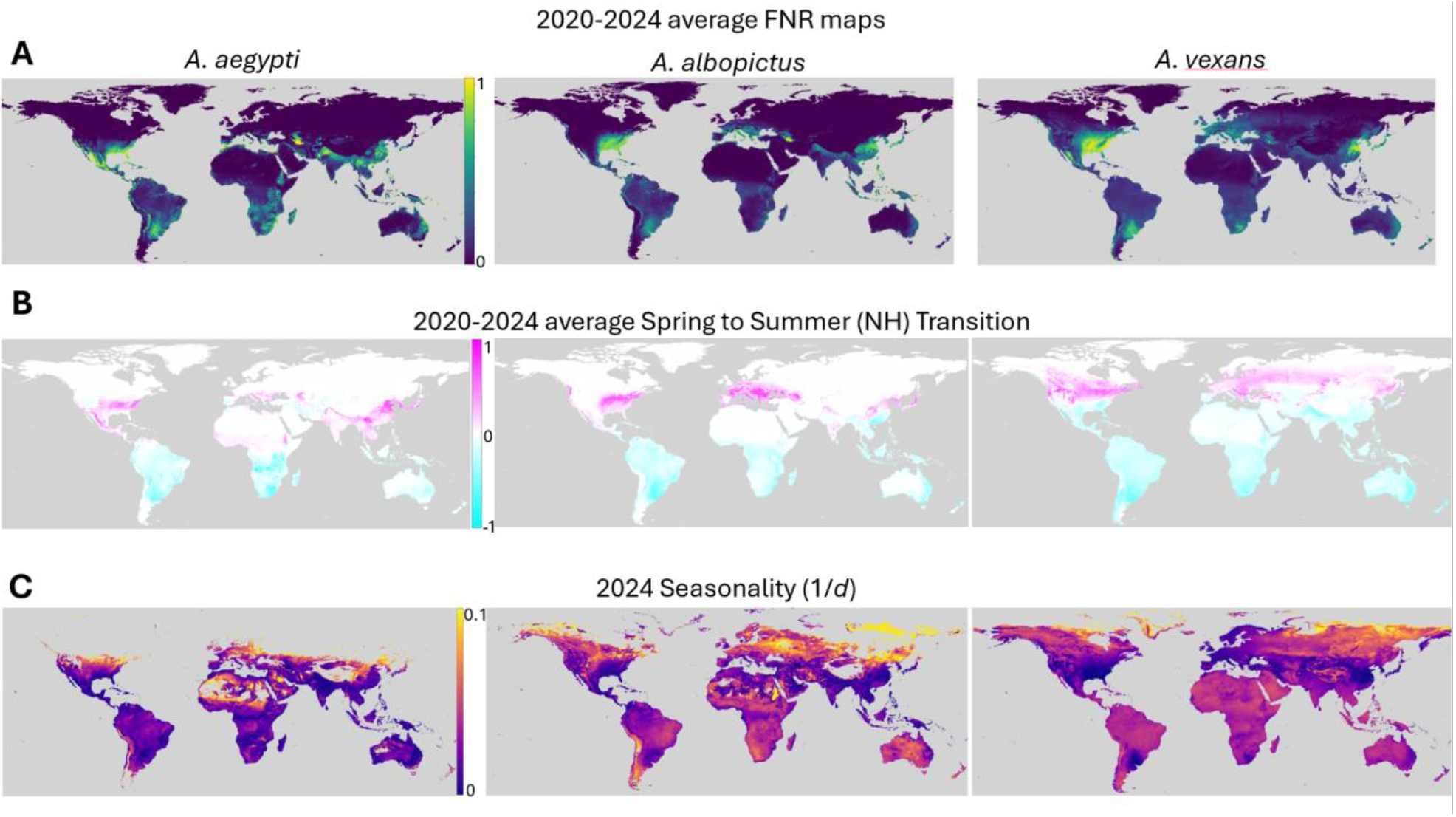
Seasonal trends. **A**. Average maps for daily FNR values in 2020-2024 for each of the three mosquito species studied. **B**. Average of differences in FNR between the northern hemisphere’s spring (March through May) and summer (June through August) for 2020-2024. **C**. Seasonality (1/*d*) where diameter *d* is the minimum number of days comprising 95% of a given location’s FNR probability mass for 2024. Color is logscale. Locations with very low cumulative abundance and consequently very short seasons as defined are assigned background color (gray), see Methods for details.

In order to evaluate the strength of seasonal transitions throughout the year and removing the limitation that seasons must begin and end on the same calendar day across the globe, we computed a seasonality score as follows. For every location, for each species, the smallest time window encompassing 95% of that location’s cumulative FNR values, *d*, was identified as that location’s “mosquito season”. We then report seasonality as *1/d* where higher seasonality corresponds to a shorter, more sharply defined, mosquito season and a lower seasonality corresponds to a longer mosquito season (Figure 3C). Generally, regions near the equator have longer, less well defined seasons than those far from the equator, as expected.

### 2.4 Historical analysis

An average map, accounting for yearly variation in short term weather patterns as shown in Figure 3A, was computed for the historical period: 1980-4. The difference map showing the change in mosquito risk over time was then computed (relative to 2020-2024), see Figure 4A. (see Figures S1, S2, and S3 for all historical windows and species). For all three species, the clear trend overall is a decrease in risk near the equator and an increase in risk in more historically temperate climates including much of the United States and Europe. For both *A. albopictus* and *vexans*, this is largely a North/South divide, while *A. aegypti* displays more significantly increased risk in South America, Southern Africa, and Southeastern Australia. Notably, major population centers in Southeast Asia are also predicted to be experiencing increased *A. albopictus* risk. We emphasize that these trends reflect an overall, marked expanse of the regions of the globe that support the success of these species. Regions at decreased risk relative to historical averages, remain among the highest risk worldwide. These trends are additionally visualized in Figures S2 and S3 which display the global change in habitability at 10 year intervals.

**Figure 4:**
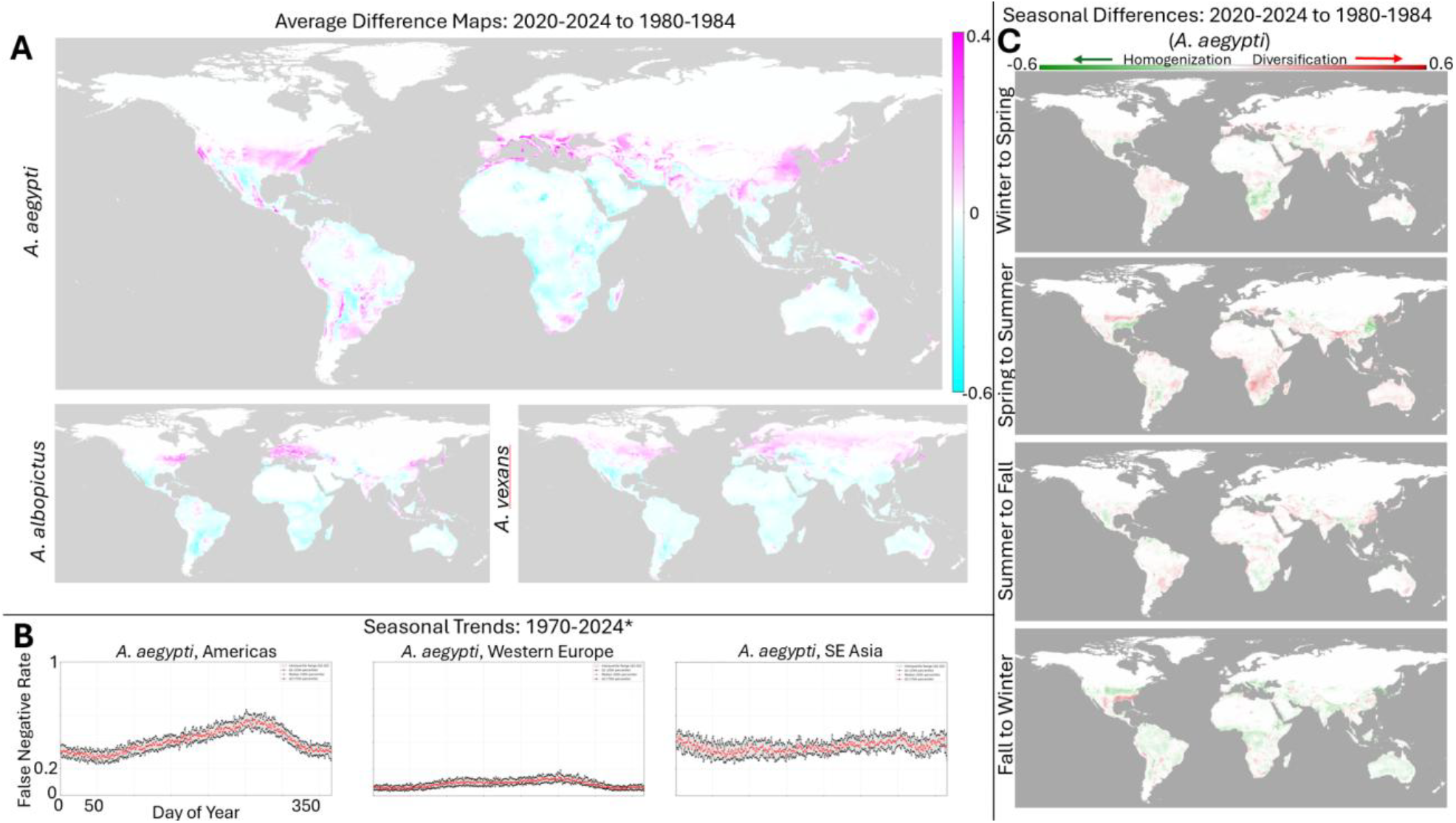
Historical trends. **A**. Global difference maps between 2020-2024 yearly average FNR and 1980-1984 yearly average FNR. **B**. Median (red), 25th percentile (black, lower) and 75th percentile (black, upper) false negative rate by day of year, from 1970-2024 (*first half-decades prior to 2010). **C**. Differences between FNR seasonal averages from 2020-2024 to 1980-1984.

Seasonal patterns have also significantly changed over the past few decades. Figure 4B displays the baseline seasonal patterns, averaged over the entire period, for the regions highlighted in Figure 2A: a strong summer-fall peak in the Americas; a weak summer-fall peak in Western Europe, and weakest seasonality observed in SE Asia (see Figure S4 for a visualizations of each year independently). Figure 4C displays the differences in the strength of seasonal transitions relative to 1980. First, seasonal transitions are mapped as displayed in Figure 3B and the historical change in the strength of these transitions is computed. For each season, negative values reflect temporal homogenization of the mosquito ecology (the difference between seasons is smaller than it has been historically) and positive values reflect temporal diversification (the difference between seasons is larger than it has been historically).

In contrast to the overall increase in mosquito range, we find that changes in seasonal patterns are more regionally variable. In particular, we highlight the change in seasonal patterns observed for *A. aegypti* within the US which has experienced winter-spring and spring-summer diversification and fall-winter homogenization. These patterns are consistent with the overall increase in mosquito habitability within the region, leading to both sharply higher peak habitability within the historical season as well as an extended duration of the mosquito season later into the year. Overall, globally, this regionally specific pattern of changes to seasonality, both diversification and homogenization, are suggestive of habitat destabilization.

### 2.5 Burden of Disease

Combining our predictions for mosquito habitability/observability at a given place and time with global population data (50) provides us with a crude estimate for the global burden of vector borne disease (Figures 5, S5). Global gridded population density was retrieved for 1990, 2000, 2010, and 2020 with a spatial resolution of 30 arc-seconds, with intermediate values estimated via linear interpolation. Similarly, daily FNR values were linearly interpolated for years where explicit model predictions were not performed. Figure 5A displays the number of person-days at high risk of observing a mosquito (FNR ≥ 0.5) of each species. Globally, *A. aegypti* population-days at-risk have increased 70% from 1990 to 2020, compared to 45% for *A. albopictus* and 26% for *A. vexans*. Due to its combination of high mosquito presence and high population, SE Asia represents a majority of global population-days at risk. While the Americas reflect global trends of *A. aegypti* as the predominant species likely to contribute to the burden of disease, western Europe’s predominant species has been *A. vexans*, with historically stable risk, followed by *A. albopictus*, with increasing risk.

**Figure 5:**
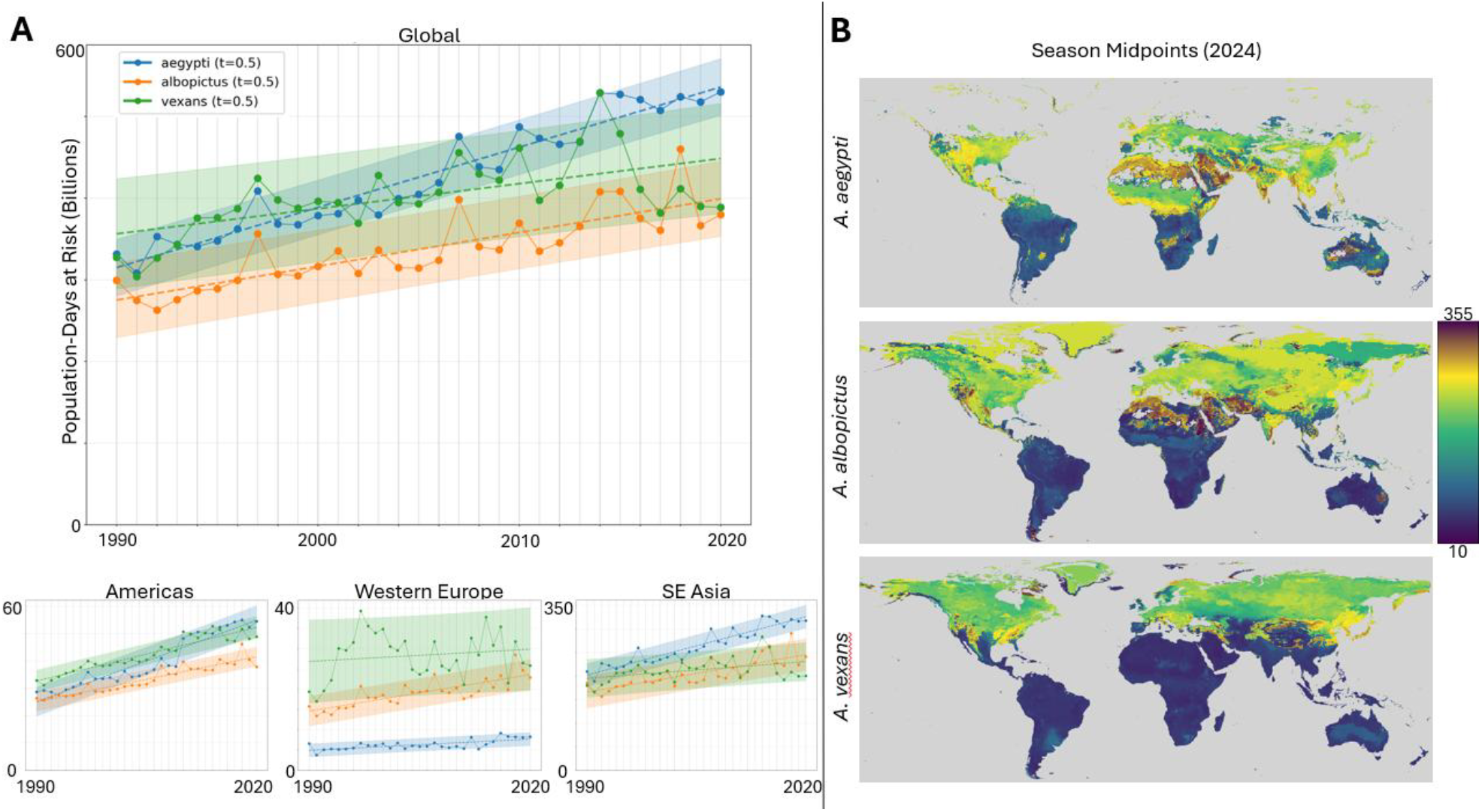
Global burden of disease by species. **A**. Population-days at risk above threshold (FNR ≥ 0.5) for each species (*A. aegypti*, blue; *A. albopictus*, orange; *A. vexans*, green) globally (top) and regionally (Americas, bottom-left; Western Europe, bottom-middle; SE Asia, bottom-right) from 1990 to 2020. Data points for each species use linearly interpolated population values between census years, and linearly interpolated FNR values for second half-decades prior to 2010. Linear regression (dashed line) shown for each species, along with 90% prediction interval (light blue, light orange, and light green shading, respectively). **B**. 2024 season midpoints, the day of the year at which roughly 50% of the cumulative FNR for that location lies before and after, for each species.

Overall, globally for all species, as well as within most regions, we observe a stable, linear increase in the risk of exposure. Our results suggest both a consistent increase in the risk of mosquito-borne disease as well as reduced predictability for when and where disease burden will be greatest, complicating mitigation efforts. Figure 5B visualizes the extent to which peak-season midpoints are both species- and region-specific.

## 3. Discussion

In this work, we characterize an ecological modelling approach supported with a lightweight feedforward neural network. Our approach produces qualitatively similar results to the popular package, MaxEnt (17), while demonstrating the potential for improved generalizability when training data are sparse. Additionally, our approach is more adaptable to a variety of input datatypes, including high-dimensional images obtained from contemporary remote sensing repositories like the ERA5 data (25) used in this work. Both model training and predictions can be achieved on a device with as little as 16GB RAM.

We used our approach to produce ∼38,000 daily predictions for relative mosquito risk for three *Aedes* species that are significant disease vectors: *A. aegypti, albopictus*, and *vexans*. Consistent with predictions that, overall, climate change is expected to increase the global burden of human pathogens (51), we find that most of the world is at a higher risk for mosquito-borne disease than it has been in the preceding decades (Figure 4A) and that, moreover, this risk is steadily, linearly increasing (Figure 5A).

These impacts are both species- and region-dependent; however, global trends show a clear expansion out from the equator to encompass more historically temperate regions. Species specificity can obfuscate a region’s overall risk, where the decrease in one species can mask an increase in another in the general public’s perception. The potential for such masking appears to be particularly strong within the Southern US and Southern Europe where *A. vexans* risk is substantially down but *A. aegypti* risk is markedly higher. The Northward shift out from the equator is particularly strong, and the world’s three largest economic centers: the United States, China, and the European Union are all projected to experience increased risk. Mosquito-borne infections currently fall into the category of Neglected Tropical Diseases, critically impacting developing, but not developed economies. This characterization is at odds with these ecological trends.

We also find evidence for the global destabilization of mosquito habitats. Seasonal trends have substantially changed over time in regionally specific ways over relatively short geographic distances (Figures 4C and 5B). These changes may make it increasingly challenging to time public health interventions. Additionally, as we have discussed in previous work, habitat destabilization resulting in the mixture of pathogen strains which were historically regionally separated, may create new evolutionary selection pressures that lead to novel adaptations with the potential to increase infectivity or virulence (52).

Our results are limited to estimating the relative risk of observing a mosquito at a given place and time based on environmental data alone. They do not take into account the population size of the vector or the geographic range of any specific pathogen of interest which may not extend to the entire range of the vector. Future work coupling these estimates with a mechanistic model for vector reproduction and migration as well as human-driven changes to the environment (e.g. deforestation) (53) open the possibility to yield quantitative estimates for the number of infections.

## 4. Methods

The FFNN was trained on ERA5 data for temperature, humidity, precipitation, and wind speed. Date-locations of sparse true-positives were retrieved from iNaturalist. Pseudo-absence selection was approximately uniformly globally distributed. Please see Figure 1A for a schematic illustrating the data pipeline.

### 4.1 Observation data - iNaturalist

Observation data was downloaded from iNaturalist (iNaturalist.org) for three mosquito species: *Aedes aegypti* (3,903 observations from March 2008 to August 2024), *Aedes albopictus* (12,615 observations from April 2002 to December 2024), and *Aedes vexans* (4,322 observations from September 2003 to December 2024). These species were chosen due to their significance as disease vectors and their high observation counts relative to other mosquito species in the iNaturalist dataset. Notably, observational data for *Anopheles* is considerably more sparse than *Aedes* and is a subject for future work.

### 4.2 Training and Input data - Climate variables (ERA5)

The European Centre for Mid-range Weather Forecasts’ (ECMWF) ERA5 reanalysis weather data was downloaded as monthly NetCDF files from the Copernicus Climate Data Store (CDS) (https://cds.climate.copernicus.eu/) and converted to daily Tagged Image File (TIF) format at a resolution of 0.25 by 0.25 degrees. Reanalysis data includes ocean locations, which we include in all calculations but exclude from visualizations. Eight weather variables were accessed: mean, maximum, and minimum 2-meter air temperature; dew point temperature; surface pressure; average precipitation; and *u*- and *v*-components of wind speed. Daily weather files were processed for all inclusive dates in 1969-1974, 1979-1984, 1989-1994, 1999-2004, and 2009-2024. If mosquito presence on a certain date is predicted using information from the prior 365 days (sequence length of 365), this allows predictions for all but the first year in each date range, i.e. the first half-decade from 1970 to 2004, and all dates in 2010 to 2024.

### 4.3 Training and Input data

Each presence observation is joined with climate data for the nearest climate grid location to that observation, for the day prior, and up to 365 days prior to the observation, providing a *sequence length* maximum of 365, which we use for the analyses presented here. This method is extendable to longer sequence lengths, but consideration must be given to memory and computation time requirements.

A pseudo-absence is generated by choosing a date randomly within the window of presence observations and a random land location weighted by the cosine of latitude to ensure an area-proportional global distribution. The pseudo-absence is rejected if its nearest land location on the ERA5 grid is shared by a presence observation. Observations and pseudo-absences are then joined with climate data at their grid location for as many days prior as are required for the specified sequence length. We then calculate total daily precipitation as average hourly precipitation multiplied by 24; temperature range as the difference of maximum and minimum 2-meter air temperature; total wind speed from the *u*- and *v*-components; and relative humidity (RH) from dew point temperature in degrees Celsius (*T*_*d*_) and mean 2-meter air temperature (*T*_*m*_) using a version of the August-Roche-Magnus equation (54) as follows: RH = 100 × (exp((17.625 × *T*_*d*_) / (243.04 + *T*_*d*_)) / exp((17.625 × *T*_*m*_) / (243.04 + *T*_*m*_))), in which exp signifies the exponential function.

### 4.4 Neural Network

Previous work has shown that prediction from weather data using neural networks can reduce computational time, increase prediction accuracy, and provide greater generalization capability compared to purely mechanistic/stochastic models (55). Here, we use a feedforward neural network (FFNN) absent any mechanistic assumptions, providing it with both weather data and mosquito observation/pseudo-absence data. The FFNN model presented here can effectively model nonlinear climate phenomena, capturing temporal dependencies through a flattened input sequence approach, without the computational overhead required by more complex architectures.

The model architecture (Figure 1B) consists of an input layer with dimension of sequence length by the number of weather variables, three fully connected hidden layers with 128 nodes, 64 nodes, and 32 nodes, respectively, and one output layer. The hidden layers use rectified linear unit (ReLU) activation. Model selection uses *k*-Fold cross-validation (here, *k* = 5) to mitigate overfitting. Training uses adaptive moment estimation (Adam optimizer, (56)) with a learning rate of 0.001. Early stopping was implemented with validation AUC as the stopping criterion, using patience values of 10 epochs during cross-validation and 100 epochs during final model training. Data preprocessing included *z*-score normalization (standard scaling) to ensure equal contribution from weather variables with varying units and scales.

### 4.5 Train-Test Splitting and Absence Ratio

The test set was constructed by randomly choosing 20% of the observation data and an equal number of pseudo-absences. The remaining presence and unused absence data were used for training the model. Multiple versions of train-test splitting were applied - Figure 1C shows the described random 80/20 split (Random), as well as an 80/20 split based on a temporal partitioning of the data where most recent 20% of observations are reserved for testing purposes (80/20). The absence ratio was set to 0.2, though absence ratios of 0.1 through 0.5 in increments of 0.1 were also considered.

### 4.6 Loss Function and Evaluation Criteria

Binary cross-entropy with logit loss was employed as the objective function, which combines sigmoid activation with cross-entropy loss for numerical stability. Area Under the receiver operating characteristic Curve (AUC) was chosen as the primary evaluation metric during training, triggering early stopping when validation AUC failed to improve.

### 4.7 Post-processing and Model Selection

Raw model probabilities at each location were converted to false negative rates (FNR) to ensure that values are well-calibrated and to provide outputs which maintain a relative ranking between locations that is insensitive to model hyperparameters (including the absence ratio). It can be interpreted as a “risk percentile”; that is, observers in date-locations with an FNR of 0.5 (50%) have a greater likelihood of making an observation than they would if they were present at 50% of date-locations where observations were made. Leave One/Two Out Analysis was conducted to assess the relative importance of individual and pairs of weather variables on model performance (Figure 1C). Sequence length was included as a variable in this analysis, with values of 30, 60, 90, and 365 days tested to determine a suitable temporal window for capturing weather-species relationships. AUC values were compared for different combinations of weather variables and sequence lengths, with special attention paid to ROC curves, as different ROC curves can produce the same AUC value.

Visual representations of seasonal diameters and midpoints (Figures 3C and 5B respectively) exclude locations with negligible cumulative FNR near the poles as follows. First locations with season midpoints occurring within 10 days of January 1st are excluded. The limits of the colorscale for Figure 3C are then set to the 2nd and 98th percentiles of the remaining locations. This heuristic procedure was established to optimize the dynamic range of the colormap for these panels and did not influence any other calculations.

## CRediT author contribution statement

EJC, KX, and NDR designed the study. EJC and KX developed computational models and ran simulations. EJC, YIW, EAK, and NDR performed analyses. HS provided computational support for the NIH HPC Biowulf cluster. EJC and NDR created the figures and wrote the paper, which was reviewed and approved by all authors.

## Acknowledgments

We thank Helen Amos, Stephanie Uz, Anna Borovikov, Michael Bosilovich, and Anthony Campbell from NASA for their insights into remote sensing dataset selection and processing. We thank members of the Evolutionary Health Group group as well as Eugene Koonin for the valuable discussions.

EJC, HS, YIW, and NDR are supported through the Intramural Research Program of the National Library of Medicine, National Institutes of Health.This work was funded by the National Institutes of Health ITCCH Program under Project # 75N98022D00019 and the City University of New York (CUNY) Graduate School of Public Health and Health Policy. Other forms of support include additional funding from the National Institutes of Health (3U01AI096299-15S1) and the CUNY Institute for Implementation Science in Population Health. This work utilized the computational resources of the NIH HPC Biowulf cluster (https://hpc.nih.gov).

Species observations were downloaded from iNaturalist, available from https://www.inaturalist.org. This work utilizes modified Copernicus Climate Change Service information. Neither the European Commission nor ECMWF is responsible for any use that may be made of the Copernicus information or data it contains. The contributions of the NIH authors are considered Works of the United States Government. The findings and conclusions presented in this paper are those of the authors and do not necessarily reflect the views of the NIH or the U.S. Department of Health and Human Services.

## SUPPLEMENTARY FILES

Movies: Yearly GIFs for 2023 & 2024, all three species

Supplementary figures:

S1: Average maps for all half-decade averages, all three species

S2: Difference maps for all half-decade averages (to 1980-4), all three species

S3: Difference maps for all half-decade averages (consecutive)

S4: Seasonality and anomaly heatmaps, all three species

S5: 3D Population overlaid with FNR difference from 2010 to 2020

## Data Availability/Code

All code required to replicate the results shown in this work is available here: https://zenodo.org/records/17041800.

