## Supplementary Figures for "Mosquito vector ecologies are destabilizing as a result of climate change"


**Authors**

Evan J. Curcio (1,2,4), Kai Xu (2,3), Harutyun Sahakyan (2), Yuri I. Wolf (2), Elizabeth A. Kelvin (1,4,5), Nash D. Rochman* (1,2,4)

**Affiliations**

1 - City University of New York Graduate School of Public Health and Health Policy, Department of Epidemiology and Biostatistics, New York, NY, United States

2 - Computational Biology Branch, Division of Intramural Research, National Library of Medicine, National Institutes of Health, Bethesda, MD, United States

3 - Courant Institute of Mathematical Sciences, New York University, New York, NY, United States

4 - Institute for Implementation Science in Population Health, City University of New York School of Public Health, New York, NY

5 - Department of Occupational Health, Epidemiology & Prevention, Donald and Barbara Zucker School of Medicine at Hofstra University/Northwell Health, Hempstead, NY, USA

Figure S1:

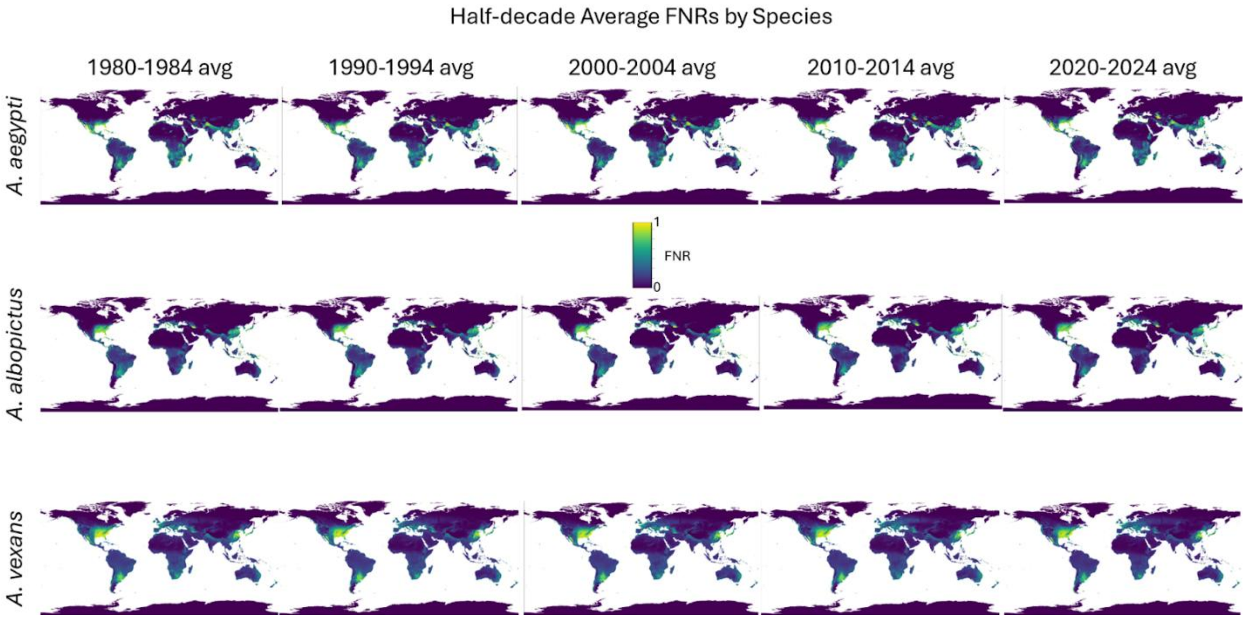


Supplementary Figure 1: Half-decade average false negative rates (FNRs) by species. Daily FNR values from each half-decade were averaged for each of the five timespans shown.

Figure S2:

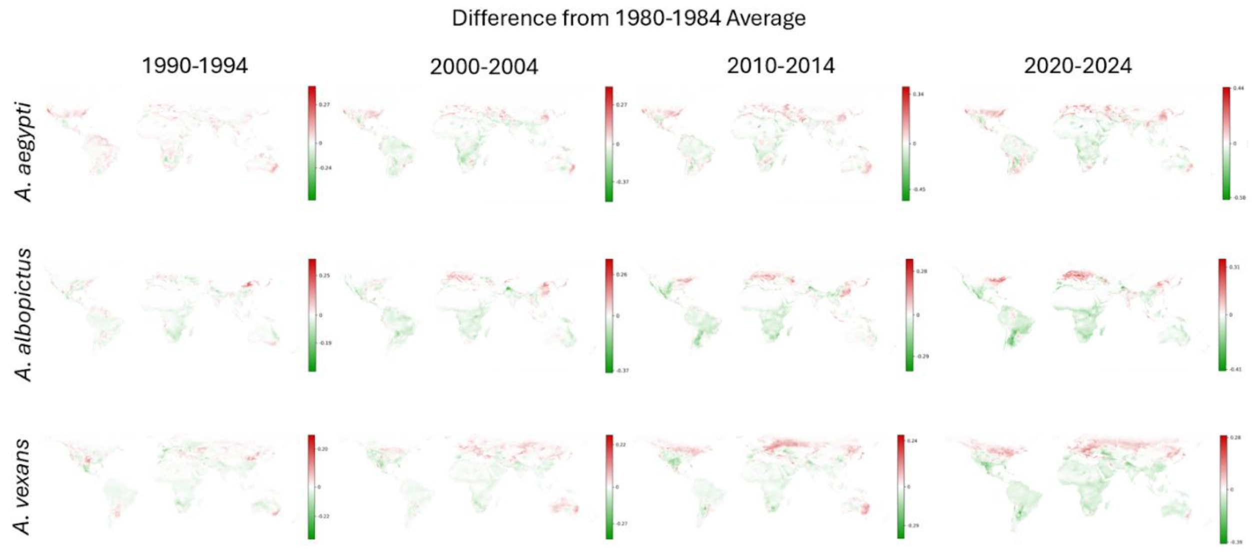


Supplementary Figure 2: Difference maps from 1980-1984 average map. Differences were taken between the 1980-1984 average value and all later timespans. Red indicates an increase in mosquito risk at a given location, while green indicates a decrease.

Figure S3:

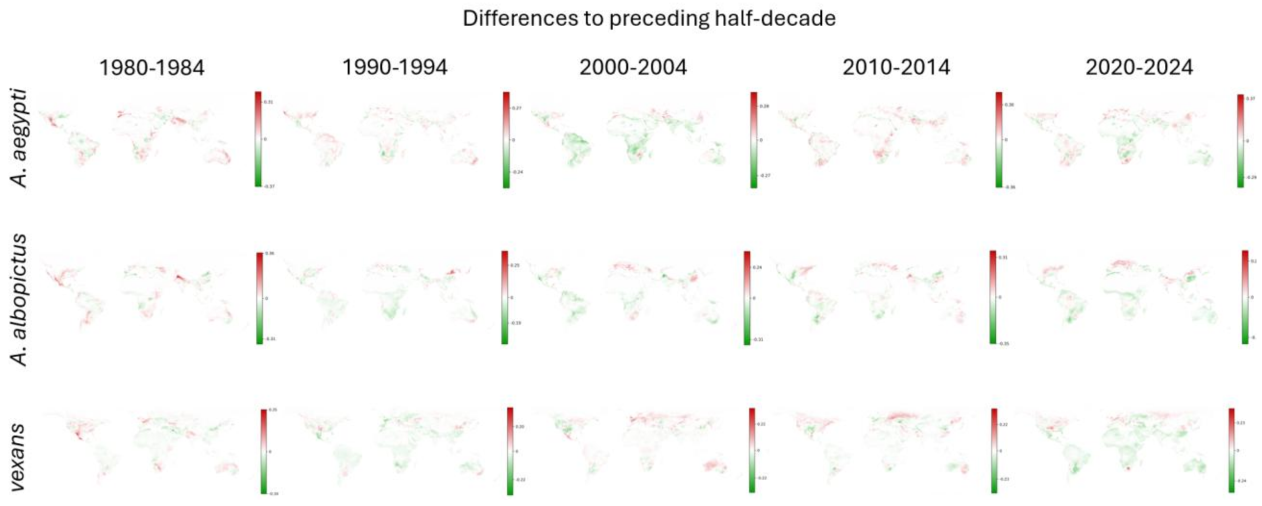


Supplementary Figure 3: Difference maps between consecutive half-decades. Differences were taken between consecutive half-decades, beginning with diff (bottom). Red indicates an increase in mosquito risk at a given location, while green indicates a decrease.

Figure S4:

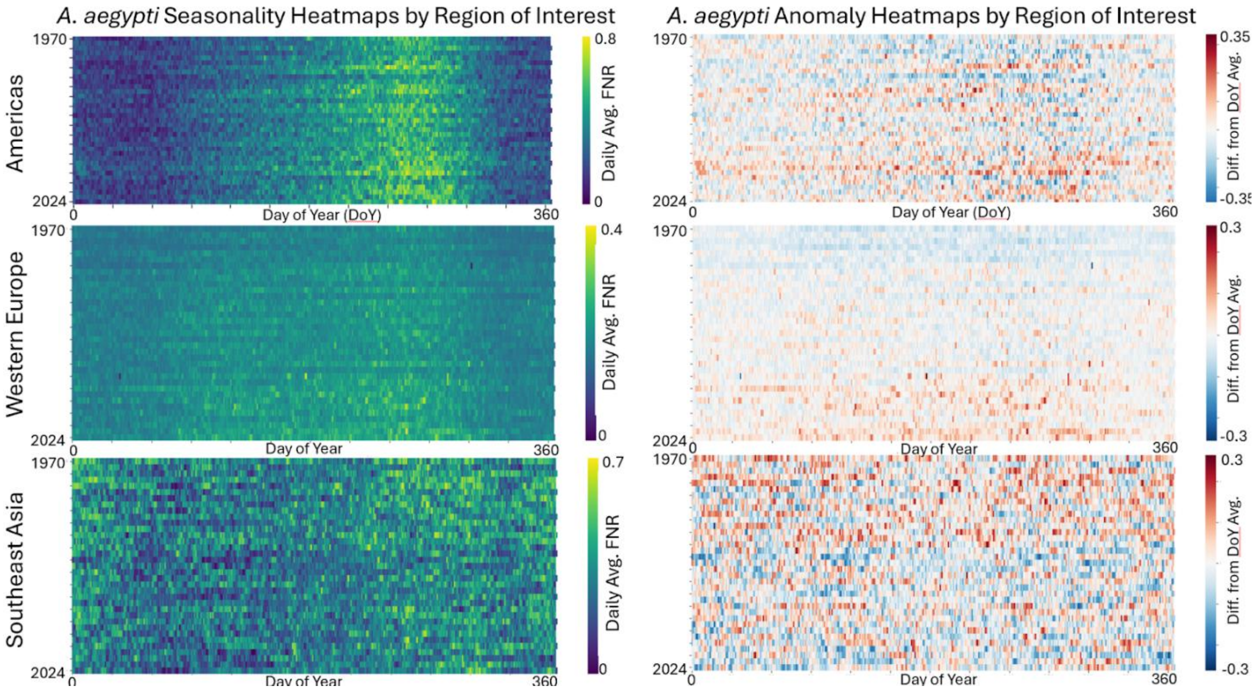


Supplementary Figure 4: Seasonality and seasonal anomaly heatmaps for *A. aegypti* by region of interest. Average FNR values are taken over the entire globe (ex-ocean) for every day of study (left). Anomaly heatmaps show the difference between a given day's average FNR value and the average across all years with the same day of year, with red indicating an increase and blue representing a decrease from the day-of-year average.

Figure S5:

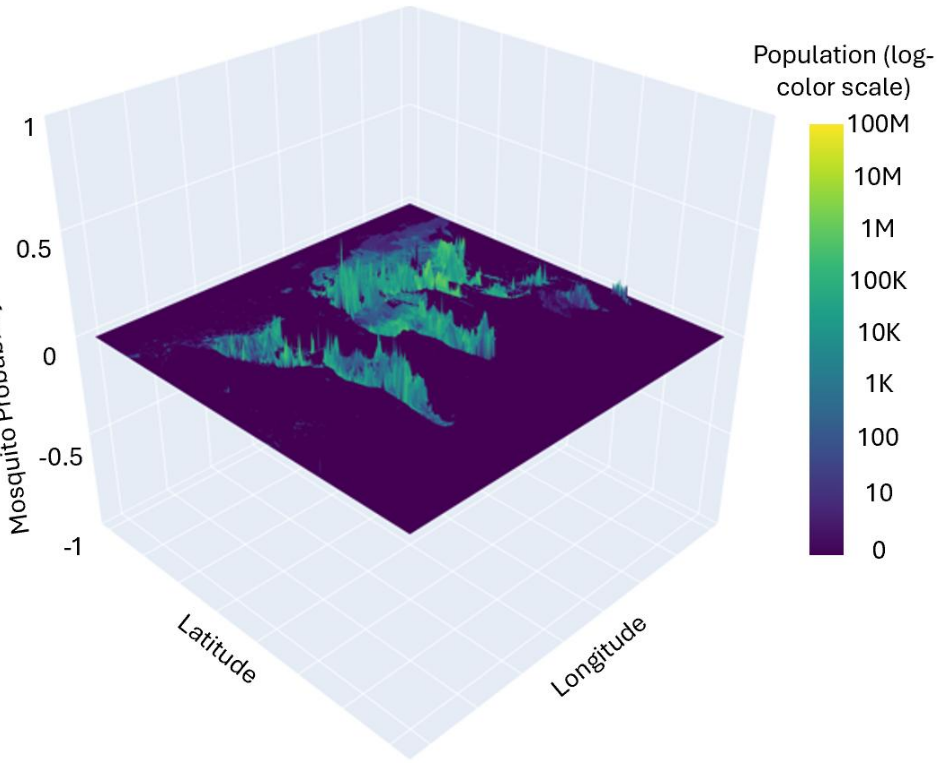


Supplementary Figure 5: 3D Plot of FNR change and corresponding population. Peaks (and troughs) represent increase (or decrease) in FNR for *A. aegypti* at a given location from 2010 to 2020. Color corresponds to the population at that location (log color scale).
