## Supplementary figures and images for "Mosquito vector ecologies are destabilizing as a result of climate change"

### aegypti_2023.gif

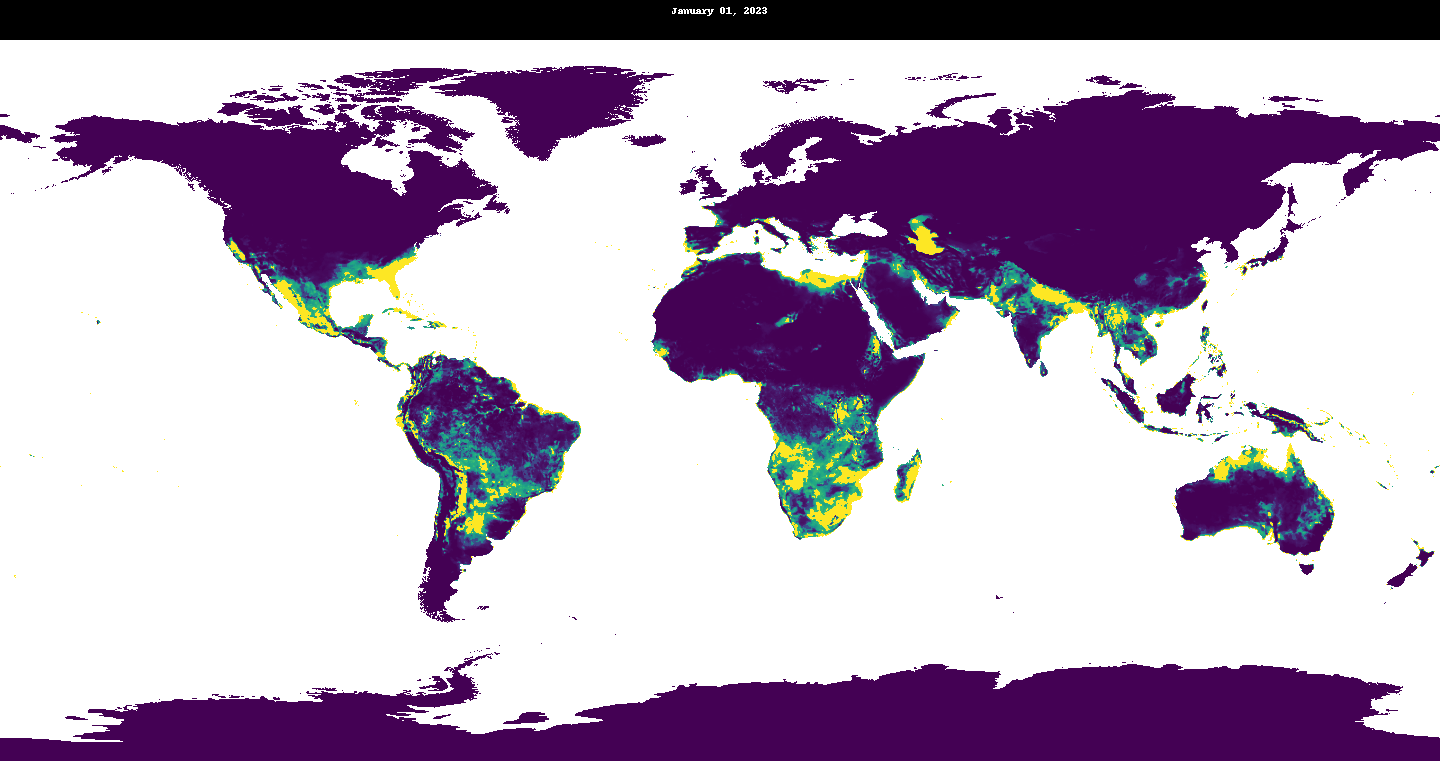

### aegypti_2024.gif

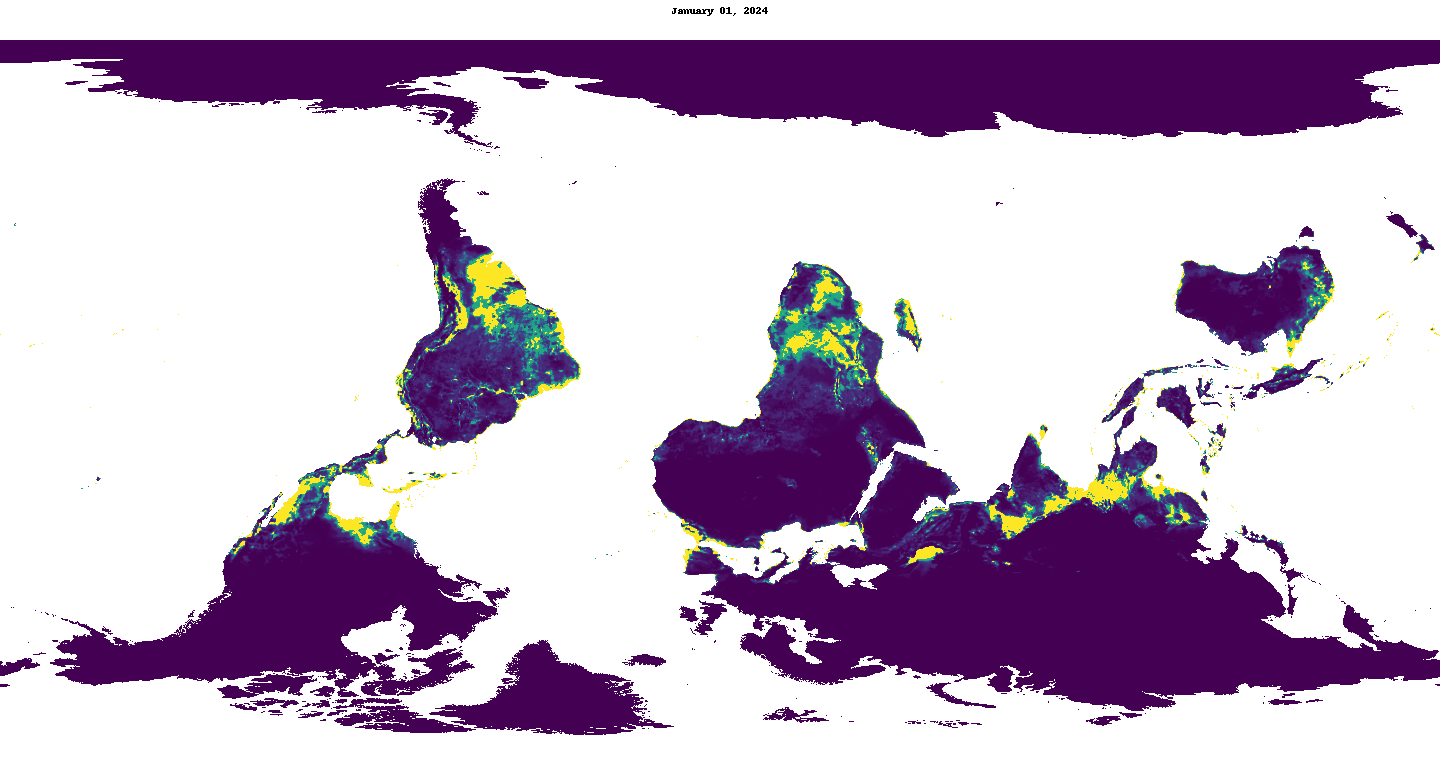

### albopictus_2023.gif

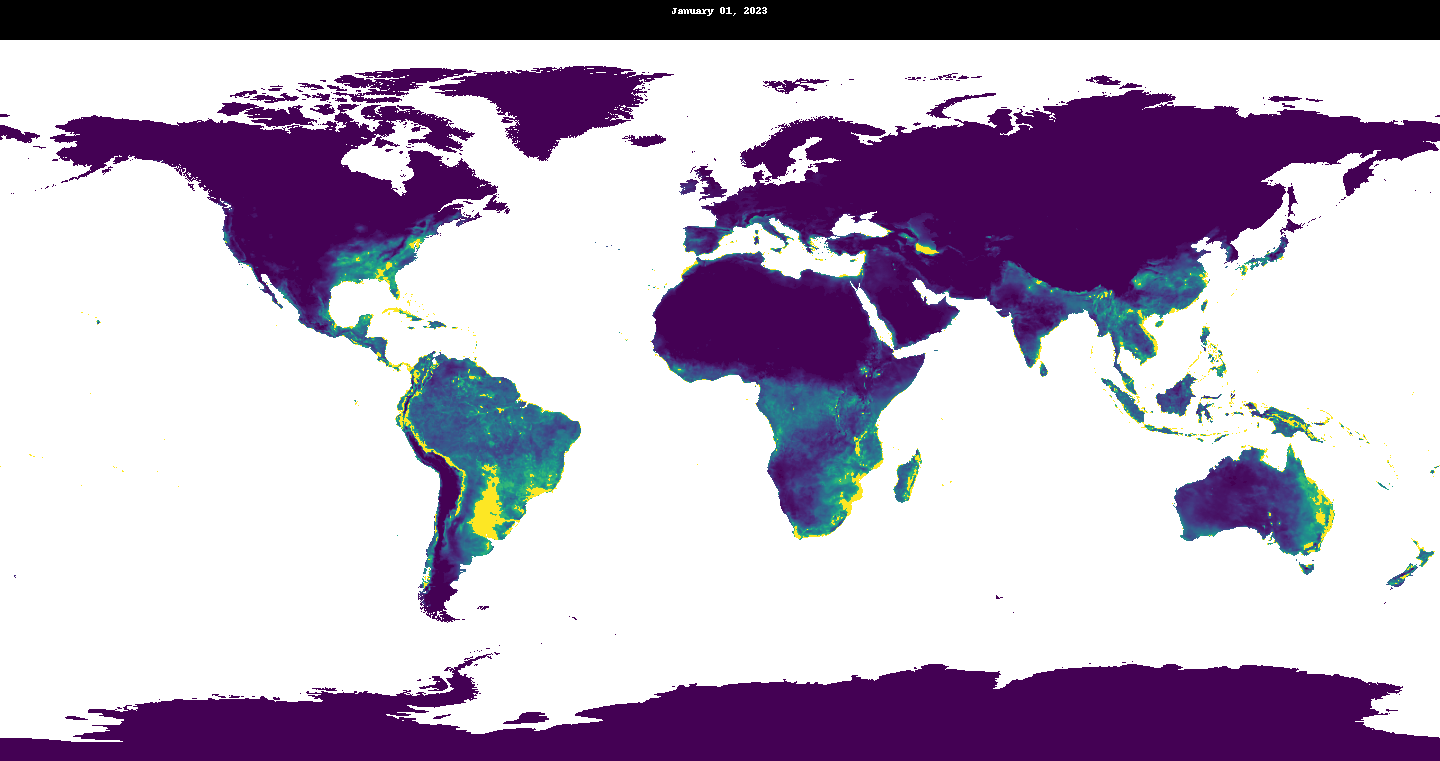

### albopictus_2024.gif

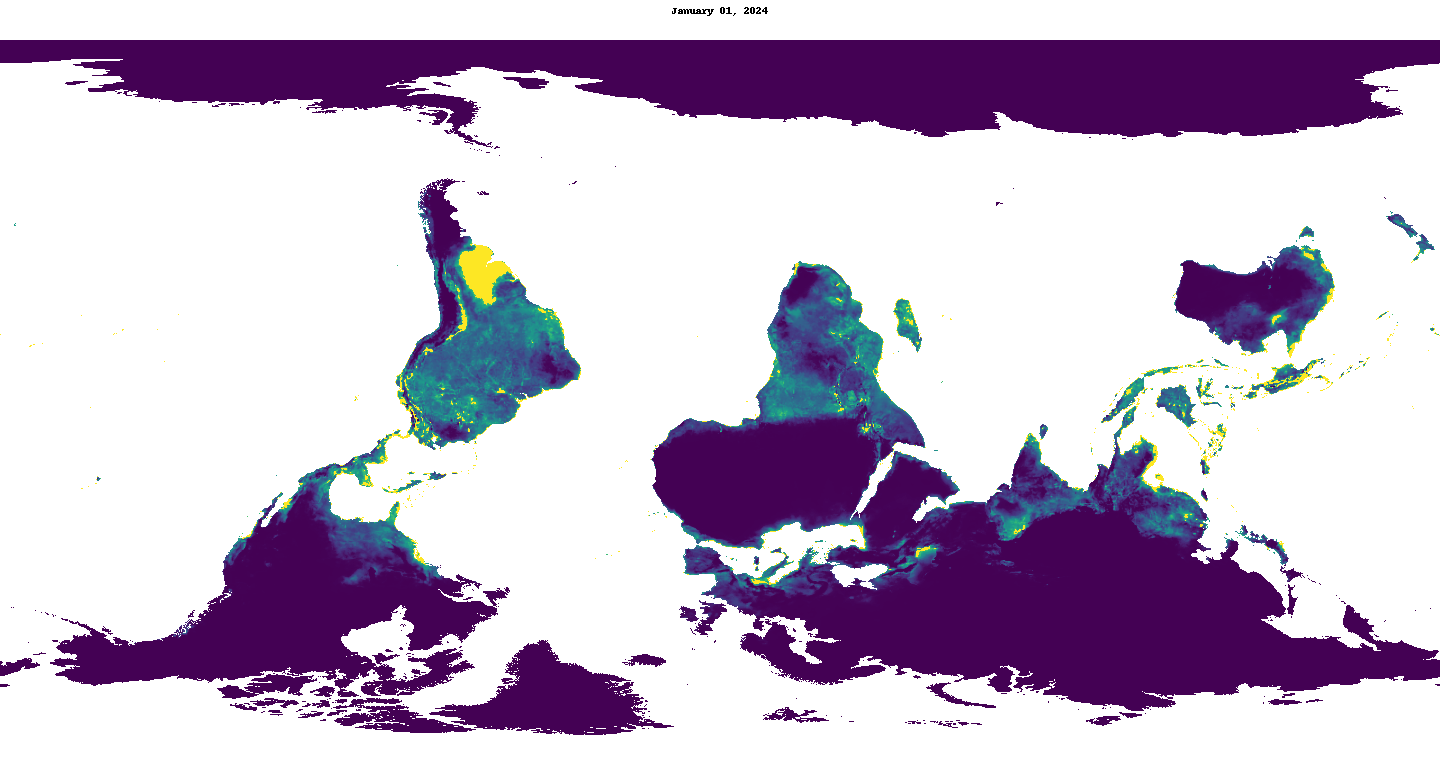

### vexans_2023.gif

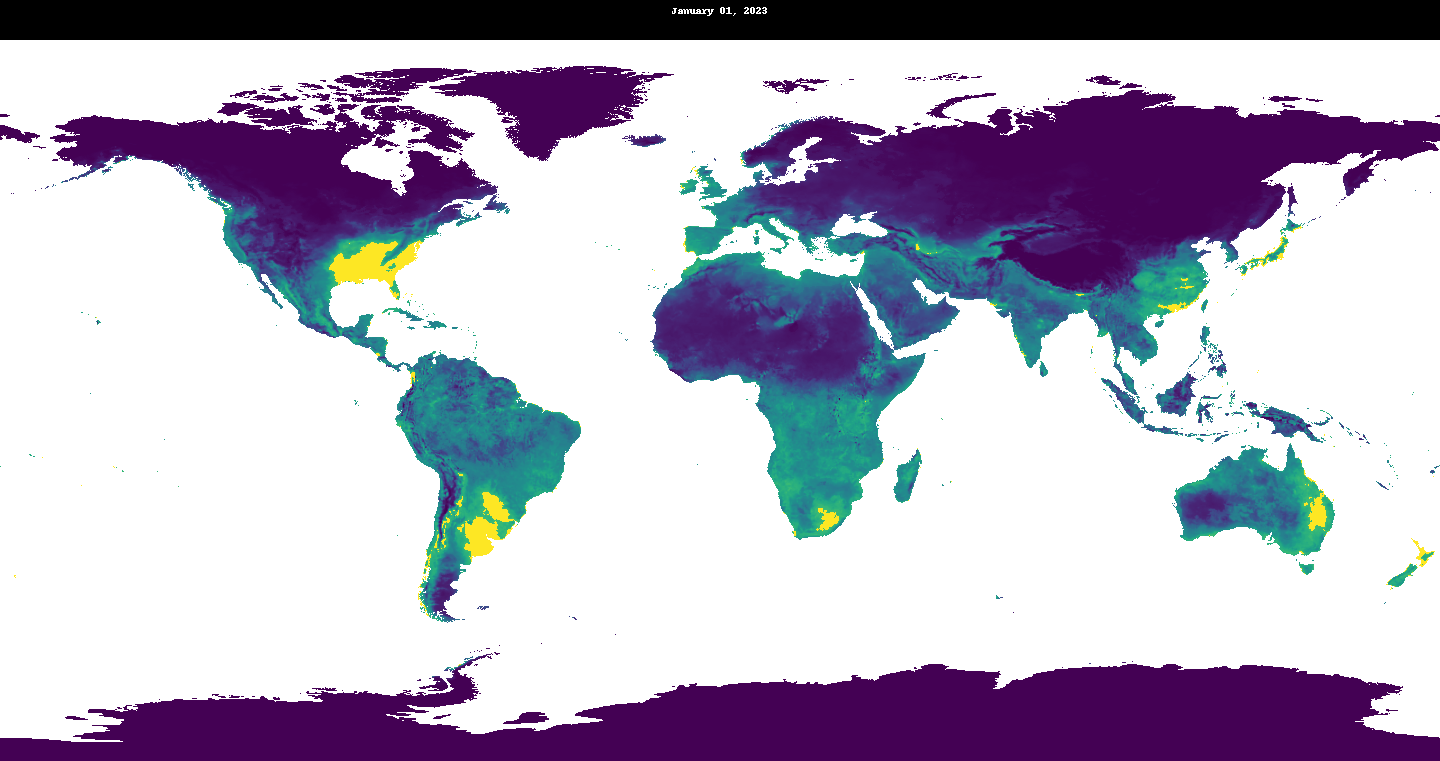

### vexans_2024.gif

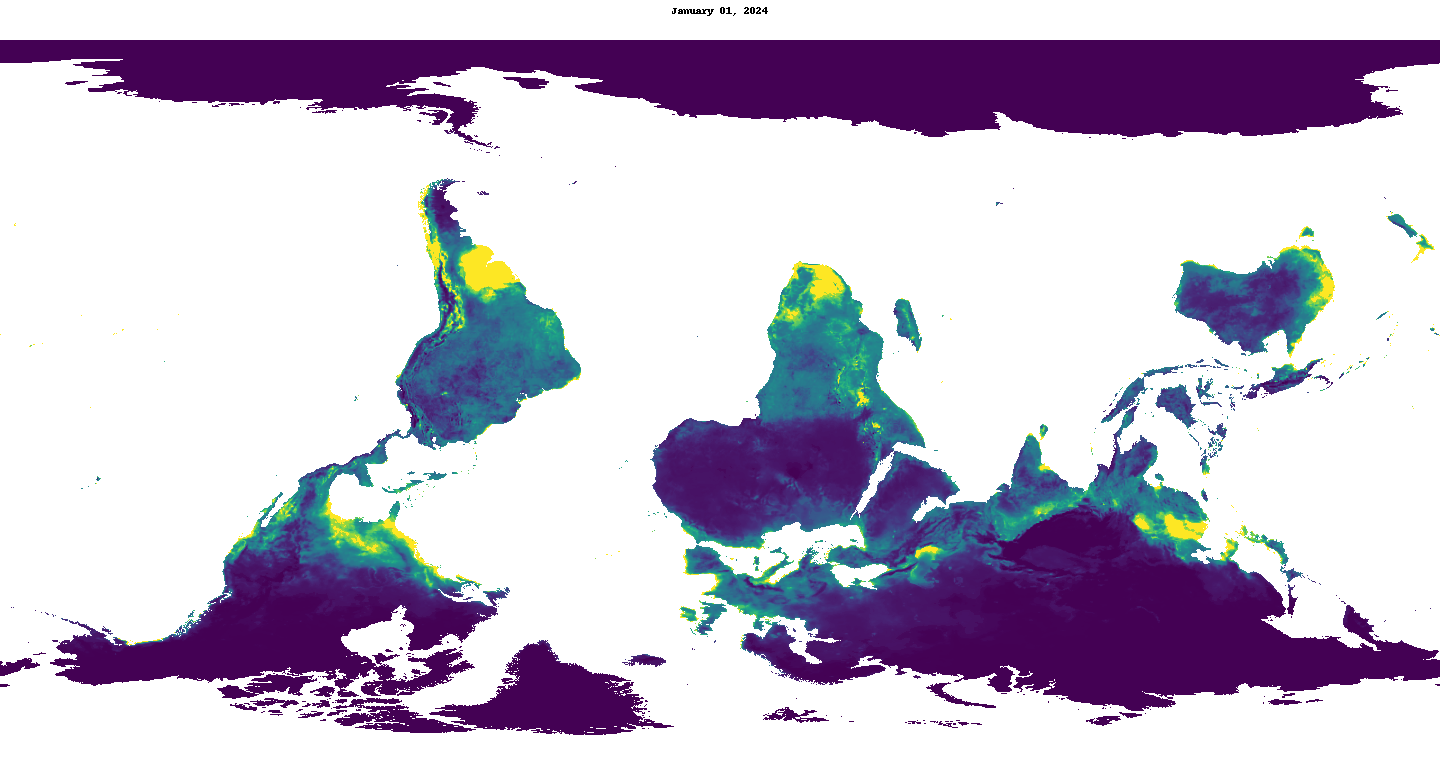
